# Automatic extraction of meaning from visual number symbols detected by frequency-tagged EEG in children

**DOI:** 10.1101/2024.03.21.585906

**Authors:** Amandine Van Rinsveld, Christine Schiltz

**Affiliations:** Laboratoire de Neuroanatomie et Neuroimagerie Translationnelles, ULB Neuroscience Institute, Université libre de Bruxelles (ULB), B-1070 Bruxelles, Belgium; FRS-FNRS Research associate, Belgium; Institute of Cognitive Science and Assessment, Department of Behavioral and Cognitive Science, University of Luxembourg, L-4366 Esch-sur-Alzette, Luxembourg

## Abstract

Symbolic number representation and manipulation is key for successful mathematical learning. However, the mechanism by which at some point in development number symbols (i.e., *1*, *2*, *3*, etc.) begin to automatically elicit useful meaning remains unresolved. Previous evidence highlighted that it is not possible to ignore the numerical magnitude when looking at number symbols, at least for adults. However, the neural mechanism behind the progressive automatization of symbol processing remains largely unknown, namely because these kinds of cognitive processes are difficult to isolate due to the general cognitive skills involved in any explicit task design. We thus developed an experimental paradigm specifically targeting the neural correlates of implicit magnitude representations by frequency-tagging magnitude changes within a visual stream of digits. Automatic magnitude processing was assessed by presenting a stream of number symbols with a frequency-tagged change of magnitude allowing to identify automatic categorization of the symbols by their magnitude in (pre)school-aged children. Stimuli were displayed with a sinusoidal contrast modulation at the frequency of 10 Hz and Steady-State Visual Evoked Potentials were recorded. These electrophysiological measurements showed a neural synchronization at the harmonics of the frequency of the magnitude changes recorded on electrodes encompassing bilateral occipitoparietal regions. The current findings indicate that magnitude is a salient semantic feature of the number symbols, which is deeply associated to digits in long-term memory across development.

**Significance Statement:** Acquiring strong semantic representations of numbers is crucial for future math achievement. However, the learning stage where magnitude information becomes automatically elicited by number symbols (i.e., Arabic digits from 1 to 9) remains unknown, namely due to the difficulty to measure unintentional automatic processing of magnitudes. We used a new experimental paradigm especially targeting the neural mechanisms involved in the automatic processing of magnitude information conveyed by number symbols. Frequency-tagged electrophysiological responses have the advantage to provide large amounts of reliable data with a high signal-to-noise ratio in a minimal amount of time. The current study is the first to take advantage of this in developmental populations to understand early automatic magnitude representations in children’s numeracy development. The electrophysiological responses demonstrate that the magnitude information is already automatically accessed from number symbols in children at the end of preschool, highlighting the importance of the first years of life for building automatized magnitude processing skills.

## Introduction

The mastery of basic mathematics is fundamental in our modern human societies, and individuals’ poor numeracy can be detrimental to various aspects of their daily life ranging from employment to well-being^1^. Indeed, large inter-individual differences exist in children’s math abilities from an early age, and they can have lifelong consequences as those differences are likely to persist into adulthood^2^. Processing the magnitude vehiculated by number symbols (e.g., *1*, *2*, *3*, etc.) is strongly linked to mathematical achievement across lifespan^3,4^. As pointed out by a recent meta-analysis, the mastery of symbolic magnitude processing is a key factor of successful mathematical learning^5^ and symbolic magnitude representations are crucial for numeracy and predict individual differences in mathematical achievement ^6 7^. However, the mechanism by which number symbols begin to automatically elicit useful meaning remains unresolved^8^. The aim of the current study is to measure semantic representation of magnitude from symbols and to better understand the developmental stage where this process becomes automatized.

Learning the meaning of number symbols occurs in parallel with a broader ongoing developmental and brain maturation course. Over the first years of life, the right intraparietal sulcus (IPS) becomes selective for non-symbolic numerosity processing ^9^. Brain regions supporting magnitude representations in infants are rather right-lateralized in the parietal cortex^10^ and become bilateral with symbolic number learning due to a left IPS specialization for symbolic numerical cognition that emerges across development^11^, though number symbols’ processing is not confined to parietal regions^12^. The emergence of symbolic numerical cognition encompasses a functional specialization of parietal regions and a shift from frontal regions, generally associated with domain-general cognition, to parietal regions when numerical operations become less effortful and more fluent across development^13^ ^14^ ^15^. Difficulties or impairments in mathematical learning have been consistently associated with weaker symbolic magnitude representations and a lack of functional specialization^16^ for a review, see^17^. However, the early onset of automatic magnitude processing from number symbols in itself is not yet understood, partly due to the difficulty to evaluate that automaticity in isolation.

Behavioral evidence converged to the idea that it is not possible to ignore the numerical magnitude when looking at number symbols, especially for numerated adults. The numerical Stroop effect (i.e., interference of the numerical magnitude when the task consists in judging the physical size, for instance: 2 vs. 5) suggests a certain level of automaticity in the processing of numbers’ magnitude, as the latter is processed even when irrelevant for the task^18^ ^19^. Moreover, neuroimaging studies showed brain activations corresponding to magnitude representation during passive viewing of number symbols^20^. This access to numbers’ magnitude seems automatic in adults but probably takes time to become established in children^21,22^. These studies suggested that the magnitude processing of the symbolic numerals becomes automatized across typical mathematical development. However, the mechanism behind the emergent automaticity of number symbols’ processing across development is still unknown because it is challenging to ensure either that the semantic of magnitude is specifically activated in passive viewing of symbols or that automatic semantic processing occurs during any explicit behavioral task.

Frequency-tagging approaches represent an opportunity to overcome the current limitations to assess the automaticity of the magnitude representation conveyed by the number symbols in behavioral or conventional experimental designs^23^. Frequency-tagging approaches consist in measuring the steady-state responses (SSR) of a population of neurons synchronizing on the frequency of an external stimulation^24^. This type of paradigms can be used to record SSR to specific aspects of processing tagged at a certain frequency of presentation, which in the present case resolves the difficulty of disentangling automatic semantic processing of magnitude from domain-general cognitive processes. Frequency-tagging has been extensively used to record low-level SSR (e.g., light flashes, click sounds) but this technique has been recently successful in reflecting higher-level order semantic processing (i.e., faces^25^, objects^26^, tools^27^, words^28^, quantities^29^). We recently succeeded to use frequency-tagging to assess the automatic processing of magnitude from symbols in an incidental manner in adults^23^. This paradigm consists in presenting Arabic digits (i.e., 1-9) periodically with a variation on the magnitude size or the parity occurring at a certain frequency, then recording the neural synchronization on the magnitude change as an index of automatic processing. This research demonstrated that adults are capable of very rapidly extracting the meaning of number symbols based on their magnitude or even their parity properties.

### Experimental paradigm

In the current study, we used a frequency-tagging EEG paradigm to measure the level of automatic processing of magnitude from number symbols in children. An important distinction needs to be made between two types of automatic processes: intentional automatic processes that are activated for the purpose of an ongoing task and unintentional automatic processes that occur even in absence of any relevance for the ongoing task^30^. Therefore, to understand the relevance of unintentional automatic access and representation of magnitude from symbols for actual number tasks performances, we assessed both the neural response recorded by the frequency-tagging paradigm and explicit magnitude judgments performed by the same individuals. The frequency-tagging paradigm consisted of fast periodic visual presentations of digits ranging from 1 to 9 at 10Hz. A periodic deviant was presented every eight stimuli (i.e., 1.25Hz). The difference between standards and deviants was either based on the magnitude (i.e., smaller or larger than 5) in the *magnitude* condition or on a random classification in the *control* condition (see Figure1). A response at the frequency of the oddball (i.e., 1.25 Hz) would mean that the system discriminates the deviants from the standards and is thus able to categorize the presented numbers based on the manipulated dimension: magnitude or random. In addition, all participants will also perform an explicit number comparison task and a number to line placement task. The comparison between the frequency-tagging responses measuring implicit processes and the outcome at more conventional explicit tasks will disentangle automatic from effortful magnitude processing that both occur when encountering number symbols and track how both types of processing interact in children.

## Results

First, we found a robust response at the base rate of presentation (i.e., 10Hz) in all conditions and groups maximal over medial occipital channels but visible all over the posterior regions (Figure 2). This simply reflects the neural response at the same frequency as the stimulation rate and shows participants were looking at the visual stream of stimuli evenly across conditions and groups. To measure the automatic discrimination of the magnitude within the streams of digits, we used the baseline-corrected amplitudes of the oddball frequency (i.e., 1.25 Hz) and summed its harmonics up to the 7th (i.e., all harmonics before the base frequency). Those sums of baseline-corrected amplitudes (SBA) were computed per participant and per condition. We found a clear response to the oddball for the magnitude condition condition, and to a lesser extent for the control condition. The strongest SBA peaks at the oddball frequency were recorded around occipito-parietal electrodes on the left (PO7, P7) and right (PO8, P8) for the magnitude condition, and mainly on the right for the control condition. To get a clearer picture of these results, we considered three posterior regions of interest (ROI) for further analyses: left occipito-parietal, right occipito-parietal, and bilaterial occipito-parietal regions (Methods).

**Figure 1.**
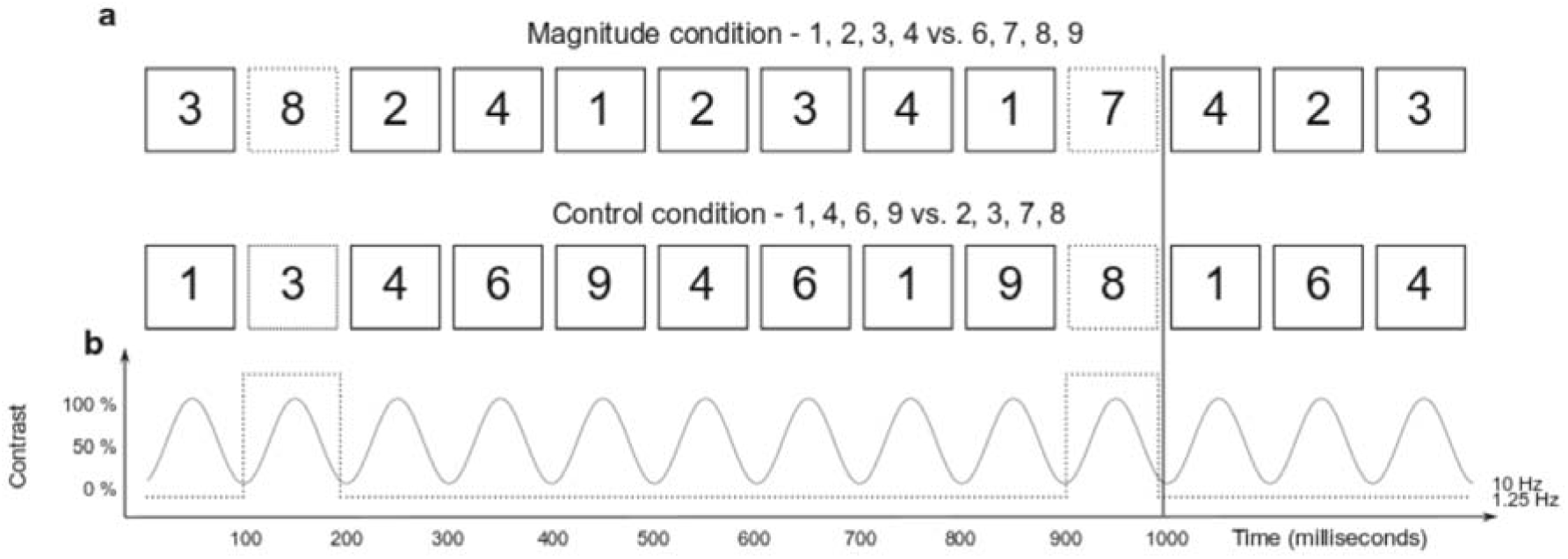
Experimental design of the frequency-tagging paradigm. Panel a depicts the stimuli presented in each condition. Number symbols are presented at a rate of 10 Hz, and a periodic deviant occurs at a rate of 1.25 Hz (i.e., one in eight stimuli). Within each sequence, numbers are be subdivided into standards and deviants based on their magnitude in the Magnitude condition, and on a random subgrouping in the Control condition. Panel b. depicts the contrast modulation used for the presentation of each stimulus.

**Figure 2.**
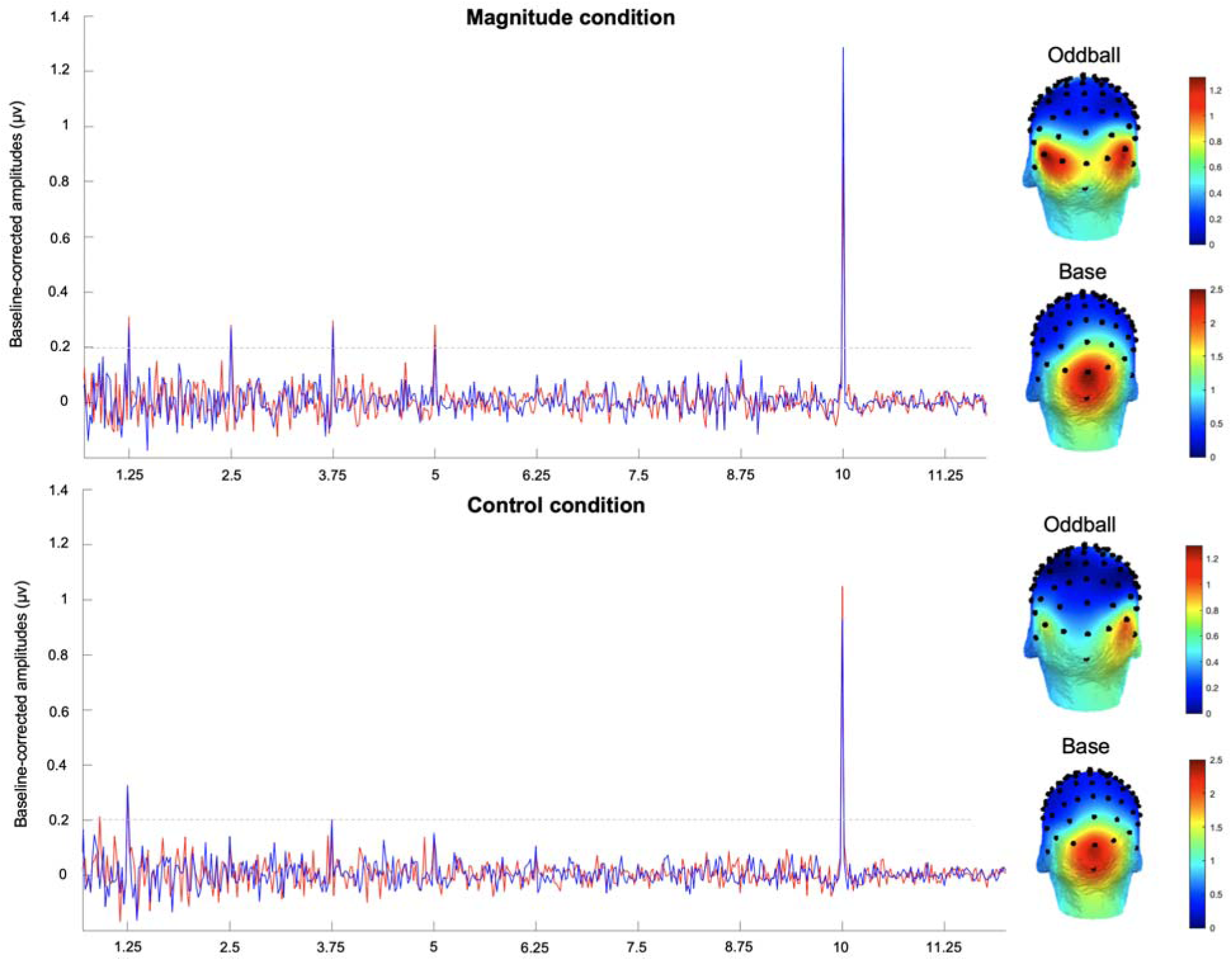
Spectrum of baseline-corrected amplitudes. Baseline-corrected amplitude (µv) as a function of frequency over the right (blue) and left (red) regions of interest. Upper panel depicts the magnitude condition and lower panel the control condition. Right panels depict the topographies of the baseline-corrected amplitudes summed over the 7 first harmonics of the oddball frequency (i.e., 1.25 Hz) and the first harmonic of the base frequency (i.e., 10 Hz). Color scale of the topographies represents the baseline-corrected amplitudes (µv).

To compare the oddball effects across conditions and groups, we ran a 2×2×2 repeated-measure analysis of variance with the condition (magnitude vs. control) and the ROI (left vs. right) as within-participant factors and the age-group as between-participant factor on the SBA as dependent variable. Results showed an effect of condition, *F* (1,30) = 4.48, *p* = .04, partial η^2^ = .13, and of group, *F* (1,30) = 12.99, *p* < .01, partial η^2^ = .30, but no effect of ROI and no interactions, *F*s < 1 and *p*s >.10 (except condition x ROI, *F* (1,30) = 1.74, *p* = .19, partial η^2^ = .05). We ran two additional 2×2 repeated measure anovas with the condition as within-participant factor and the group as between-participant factor, one for each ROI. On the left, we found an effect of condition *F* (1,30) = 6.73, *p* = .01, partial η^2^ = .18 and of group, *F* (1,30) = 12.20, *p* < .01, partial η^2^ = .29. On the right, only group effect reached significance, *F* (1,30) = 5.79, *p* = .02, partial η^2^ = .16, but no effect of condition or interaction was observed, *F*s < 2 and *p*s >.10 . Oddball responses thus seem more important for the magnitude than for the control condition prominently on the left side. This tendency can be observed in both groups from the separate scalp topographies of the SBA at the oddball frequencies as depicted in Figure 3.

**Figure 3.**
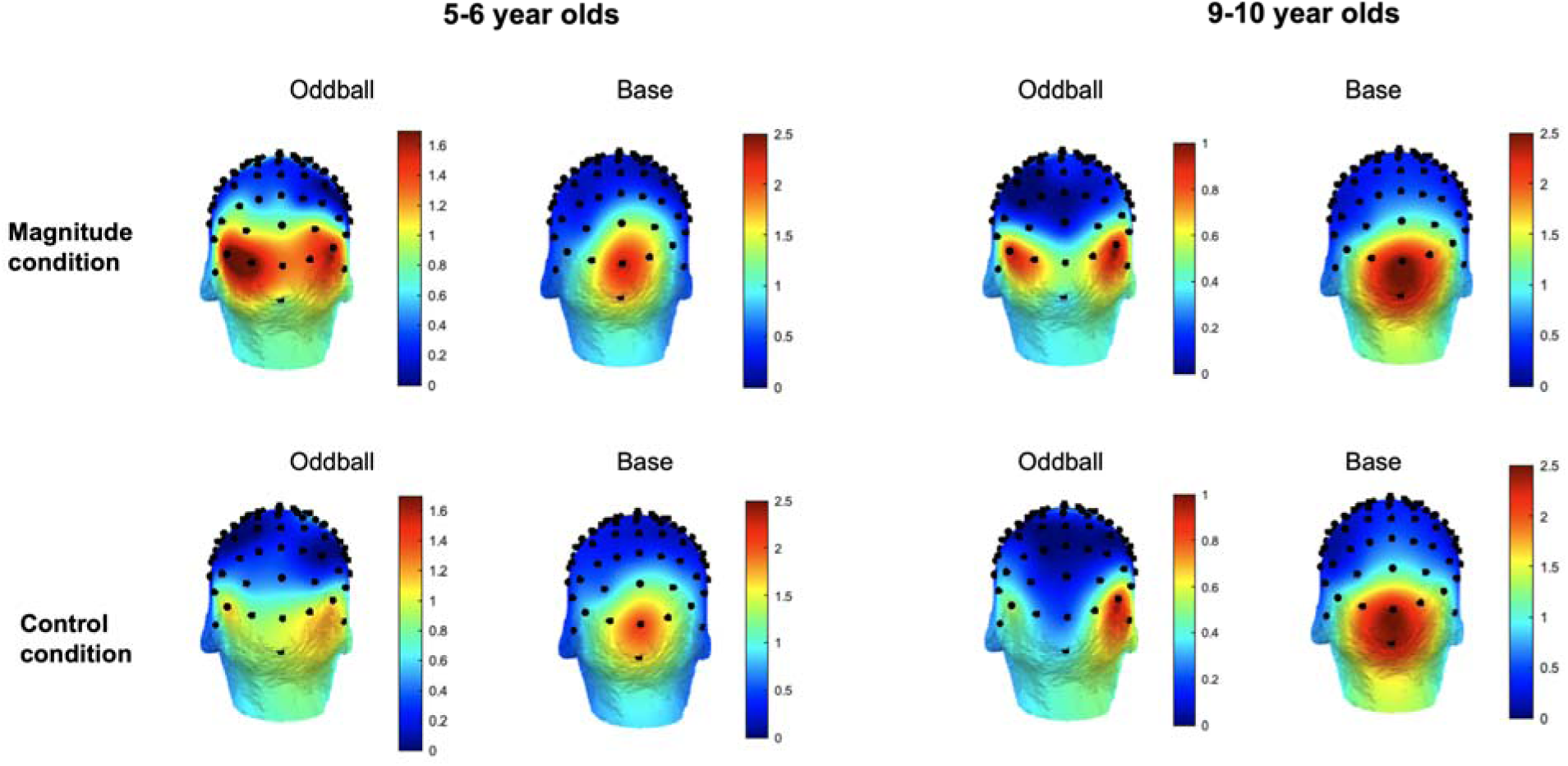
Sum of baseline-corrected amplitudes per age group. Topographies of the oddball and base responses in each group for each condition. The baseline-corrected amplitudes summed over the 7 first harmonics of the oddball frequency (i.e., 1.25 Hz) and the first harmonic of the base frequency (i.e., 10 Hz). Color scale of the topographies represents the baseline-corrected amplitudes (µv).

We further computed the z-scores to assess the presence of oddball responses in each condition against zero. We found the most compelling evidence of oddball responses when looking at the bilateral responses gathering together left and right ROIs, showing clear above threshold responses (Z > 1.64) specifically for the magnitude condition in all groups (Figure 4).

**Figure 4.**
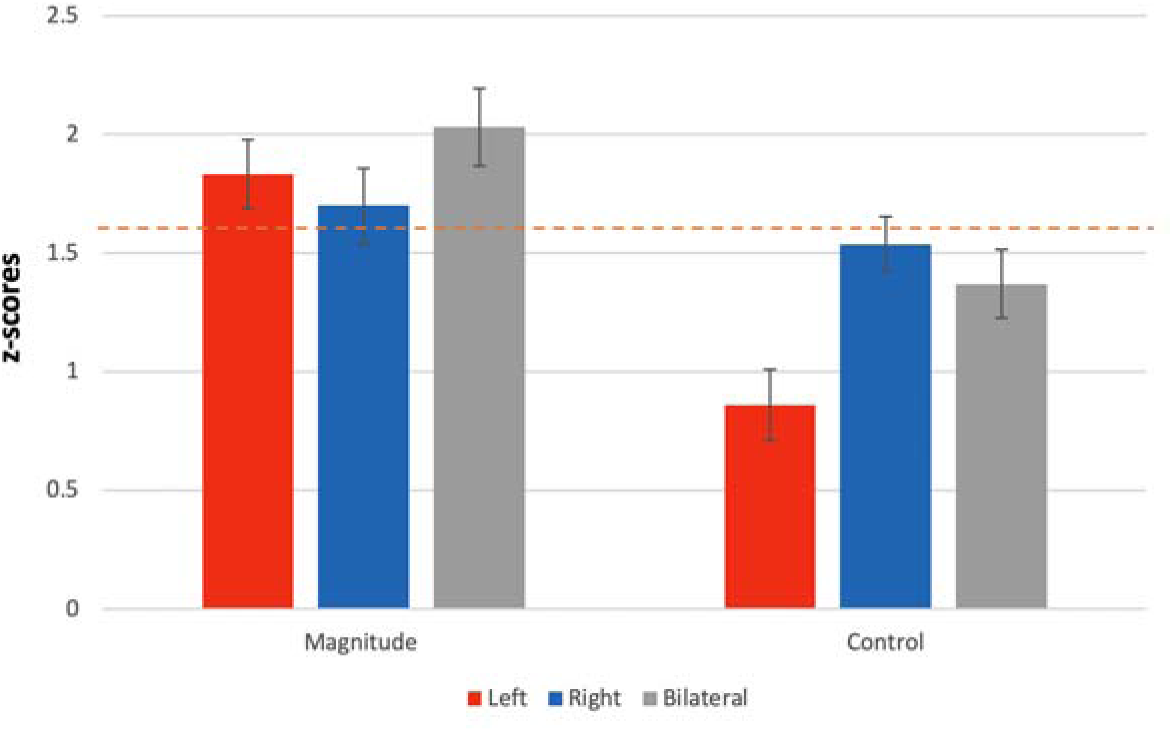
Scalp distribution of the oddball responses in z-scores. Colors depict the regions of interest: Left occipito-parietal, right occipito-parietal, and bilateral. Error bars denote the standard error from the mean. Horizontal dashed line shows the threshold of significance of 1.64.

Finally, we computed correlations to assess the link between the electrophysiological responses to the magnitude and control category detection and the behavioral measures of numerical skills (Table 1). Interestingly, we found a correlation between the behavioral number comparison and the bilateral oddball responses of the magnitude condition, r = .353, *p* = .045 but clearly not the ones of the control condition, r = -.007, *p* = .970. Other behavioral metrics of number representation on a number line did not correlate with electrophysiological measures, showing the specificity of the link between magnitude judgments at the behavioral and electrophysiological levels.

**Table 1.** Correlations between oddball responses and behavioral measures of numerical skills.

| Spearman's Correlations |  | 1. | 2. | 3. | 4. | 5. | 6. | 7. | 8. | 9. |
| --- | --- | --- | --- | --- | --- | --- | --- | --- | --- | --- |
| 1. Magnitude L | Spearman's rho | — |  |  |  |  |  |  |  |  |
|  | p-value | — |  |  |  |  |  |  |  |  |
| 2. Magnitude R | Spearman's rho | 0.186 | — |  |  |  |  |  |  |  |
|  | p-value | 0.298 | — |  |  |  |  |  |  |  |
| 3. Magnitude bilateral | Spearman's rho | 0.735*** | 0.769*** | — |  |  |  |  |  |  |
|  | p-value | < .001 | < .001 | — |  |  |  |  |  |  |
| 4. Control L | Spearman's rho | -0.020 | -0.163 | -0.063 | — |  |  |  |  |  |
|  | p-value | 0.915 | 0.372 | 0.732 | — |  |  |  |  |  |
| 5. Control R | Spearman's rho | 0.186 | 0.034 | 0.124 | 0.549** | — |  |  |  |  |
|  | p-value | 0.306 | 0.855 | 0.496 | 0.001 | — |  |  |  |  |
| 6. Control bilat | Spearman's rho | 0.121 | -0.123 | 0.025 | 0.921*** | 0.809*** | — |  |  |  |
|  | p-value | 0.508 | 0.502 | 0.891 | < .001 | < .001 | — |  |  |  |
| 7. Number comparison | Spearman's rho | 0.275 | 0.185 | 0.353* | 0.059 | -0.043 | -0.007 | — |  |  |
|  | p-value | 0.122 | 0.302 | 0.045 | 0.746 | 0.814 | 0.970 | — |  |  |
| 8. Number to line 0-20 | Spearman's rho | 0.255 | 0.142 | 0.240 | 0.236 | 0.282 | 0.243 | 0.753*** | — |  |
|  | p-value | 0.152 | 0.430 | 0.178 | 0.194 | 0.118 | 0.180 | < .001 | — |  |
| 9. Number to line 0-100 | Spearman's rho | 0.182 | 0.063 | 0.167 | 0.245 | 0.205 | 0.245 | 0.707*** | 0.725*** | — |
|  | p-value | 0.311 | 0.728 | 0.353 | 0.177 | 0.259 | 0.176 | < .001 | < .001 | — |

\* $p < .05$ , \*\* $p < .01$ , \*\*\* $p < .001$

## Discussion

The objective of the current study was to assess the emergence of automatic access to magnitude information conveyed by number symbols in an incidental manner in children. We observed frequency-tagged responses corresponding to the oddball, which consisted in a magnitude change within a stream of visually presented number digits ranging from 1 to 9. These electrophysiological responses suggest an early automatized access to the meaning of number symbols in children at the early stages of mathematical learning. Although, the quality of the magnitude representation conveyed by the number symbols is difficult to assess. Moreover, domain-general cognitive processes represent confounding factors associated with explicit magnitude judgment tasks. The frequency-tagging paradigm of the current study avoids those confounds and demonstrates the automatic nature of symbolic numbers’ magnitude representation in children.

Interestingly, we observed no differences between groups in the oddball responses, suggesting that children are already able to automatically extract magnitude information from symbols at the end of preschool. This is consistent with previous behavioral evidence showing numerical-Stroop effects at the end of preschool^31^ and cross-sectional comparisons showing no fundamental change of Arabic number processing between 2^nd^ and 4^th^ grade^19^. However, previous evidence was typically obtained from numerical-Stroop tasks where the observed effects of interferences of magnitude on physical size judgments could be resulting from inhibition or general cognitive mechanisms and not (only) reflect automatic processing of magnitude information from symbols. The current results suggest that even purely unintentional and automatic processing of the magnitude information from 1 to 9 symbols starts relatively early in mathematical learning and is indexed objectively by electrophysiological responses. We show that this type of processing seems already well established around the start of formal schooling, which is a critical time point of children’s number learning^3,32^, and may extend to associating semantic meaning to symbols in general.

We observed frequency-tagging oddball responses to the magnitude changes but also the control condition elicited some kind of oddball responses similarly as previously reported in adults^23^, suggesting some sensitivity to any type of repeated categorical grouping of the digits with the visual stream.

However, those responses were weaker than the magnitude responses and right-lateralized similarly to adults. Moreover, only magnitude responses and not control ones correlated with behavioral performances to judge the magnitude of digits. Together, those evidence suggest that even if some categorical learning may occur, it is not the sole mechanism at play explaining the oddball responses to the magnitude changes within the visual sequences.

Previous studies have demonstrated the automatic extraction of magnitude information from symbols in adults with frequency-tagging EEG centered around the right posterior sensors^23^ ^33^, while in the present children study, the oddball responses were balanced bilaterally. One previous study found left lateralized SSR by when using number words and a combination of number words and digits in adults^34^. It is possible that children who learned number symbols more recently than adults still activate left-lateralized regions along with right-lateralized activations when accessing semantic representation of number symbols due to a systematic concurrent activation of their verbal form^35^. Another possibility is that their functional specialization of the parietal regions for symbolic number semantic content is still ongoing. Indeed, previous evidence has shown that left IPS gets specialized for number symbols during development and undergoes dynamic functional changes related to the learning of numbers, while the right IPS function remains more stable across development^36^. Going into the same direction, another study comparing numbers presented within streams of letters elicited specifically right-hemisphere responses in first graders^37^.

Behavioral measures of magnitude comparisons correlated with bilateral automatic magnitude processing but not with the control condition. Moreover, despite the strong correlation between both behavioral tasks: magnitude comparison task and the placement of numbers on a number line, only the former but not the latter was correlated to electrophysiological metrics, showing the specificity of the symbolic magnitude processing targeted by the frequency-tagging oddball paradigm. Acquiring strong symbolic numeracy skills is important for future math achievement^38^ and the present experiment represents a way to assess those responses at the electrophysiological level avoiding the usual confounds of existing behavioral and electrophysiological paradigms. Together, our results suggest that unintentional automatic processing of magnitude from symbols is an early milestone in numeracy.

## Conclusion

The current study capitalized on an experimental frequency-tagging design to measure automatic brain responses to semantic changes in the magnitude of fast visual presentation of digits across development. The results showed that posterior regions can detect change in the magnitude of number symbols in a fast visual stream. Children can discriminate magnitude automatically through number symbols before entering elementary school. Future research would be needed to target individual differences in the emergence of those automatic processes across typical and atypical development.

## Methods

### Participants

Thirty-three children participated in the study. Twelve children were between 5 and 6 year old (mean age = 5 years and 5 months) and twenty-one children were between 9 to 10 year old (mean age 10 years and 6 months). All participants were French native speakers and always had French as instruction language at school. Participants of the younger group were attending the last year of kindergarten, and participants of the older group were in their fourth year of elementary school. We recruited participants with the following inclusion criteria: right-handed, normal or corrected vision, no history of learning impairments, no history of psychiatric or neurological disorders, no known difficulties with mathematical learning. We determined the sample size by considering the effect size of 0.18 as reported by Poncin, Guillaume et al. (2021) for frequency-tagged responses to digit magnitude changes in adults within the posterior regions. With an alpha of 0.05 and a power of 0.95, a sample size of n=17 should be sufficient to detect those type of brain responses. We followed APA ethical standards in the administration of the study protocol, which were approved by the Ethical Committee of the Faculty of Psychology (Study n°016/2019). Participant’s oral consent and an authorization of their legal representative were obtained after reading the information in an age-appropriate language prior to the start of the study. Participants received a gift voucher of 20€ for their participation.

### Fast Periodic Visual Stimulation

We used the FPVS variations of the oddball design where digits were flashing at 10Hz, and we introduced a periodic fluctuation within the sequence: the stimulus category changed every eight items (*i.e.*, at 1.25 Hz) and the change is referred to as the oddball. Two different conditions each consisted in a different category change^23^. In the *magnitude condition*, periodical variations were based on the magnitude of the digits (*i.e.*, smaller than 5 or larger than 5). The greatest numbers were standards and the smallest ones were deviants. In the *control condition, there were* two arbitrary categories with equal amounts of odd, even, small and large numbers (1, 4, 6, 9 vs. 2, 3, 7, 8). Digits were sequentially presented at the fast pace of 10 stimuli per second (i.e., 10 Hz, the base frequency) following a sinusoidal contrast modulation from 0 to 100%^39^(see Fig. 1). Following previous recommendations^40^, and to decrease habituation to the visual properties of the stimuli, we introduced physical variations within the stimuli stream. The stimulus font randomly varied among four possibilities (Arial, Times New Roman, Cambria, and Calibri), and the position of the stimulus randomly varied on both the vertical and horizontal axes (with a variation of maximum 10% from the center of the screen), and the font size varied (from 122 to 148, with an average value of 135). These random visual variations occurred at each onset and were thus not congruent with our frequencies of interest (*i.e.*, harmonics of 1.25 Hz). We used MATLAB (The MathWorks) with the Psychophysics Toolbox to display the stimuli and record behavioral data^41^ ^42^. The electroencephalography (EEG) recording took place in a shielded Faraday cage (275cm×195 cm×280 cm). Participants were comfortably seated at 60cm from a screen (24-inch LED monitor, 100Hz refresh rate, resolution of 1,024×768pixels), with instruction to keep their gaze at the center of the screen on a fixation point. The stimuli were displayed with a horizontal visual angle of 5° and in the center of the screen on a grey background.

### Procedure

Although classical FPVS paradigms only involve passive viewing of the stimuli, we introduced a basic orthogonal active task during the recording sessions to ensure that participants were looking at the center of the screen. Participants were instructed to fixate a small blue diamond (12px size) located at the center of the screen and to press a button with their right forefinger when they detected that the diamond changed from blue to red color. This change was not periodic and could randomly occur six to eight times in a sequence. Participants were also informed that black digits ranging from 1 to 9 – excluding 5 – would quickly appear on the screen. Each sequence was lasting 48 seconds including a gradual fade-in and fade-out of 2 seconds at the beginning and at the end, which were not included in the analyses. The sequences of each condition were repeated four times in a row and order of condition was balanced across participants. If visual inspection of the signal during acquisition led to detecting obvious offsets of the electrodes (e.g., movement), the experimenter excluded and reran the noisy sequence. This procedure occurred in less than 2% of the sequences. Before running the EEG session, participants were administrated behavioral tasks: paper-pencil number to line placement task^43^, and a number comparison task on computer. For the number to lines, participants were instructed to place a mark corresponding to the number displayed at the top of the page. We presented ten number lines with numbers ranging from 1 to 20 and ten from 1 to 100. For the number comparison task, two digits appeared simultaneously on the screen and participants were instructed to press a button corresponding to the largest of both numbers. Number pairs remained on the screen until participants responded. A total of 76 number pairs were presented to each participant in a randomized order. The visit lasted about two hours in total.

### EEG recording

EEG data were acquired at 1,024 Hz of sampling rate using a 64-channel BioSemiActiveTwo system (BioSemi B. V.). The electrodes were positioned according the standard 10–20 system locations (for exact position coordinates, see http://www.biosemi.com ). Two additional electrodes, the Common Mode Sense (CMS) active electrode and the Driven Right Leg passive electrode, were respectively used as reference and ground electrodes. Offsets of the electrodes, referenced to the CMS, were held below 40 mV.

### Data analyses

Preprocessing and analyses of the EEG data were conducted with Letswave 7 (https://github.com/NOCIONS/letswave7). Data files were downsampled from 1,024Hz to 512 Hz. Data were filtered with a four-order band-pass Butterworth filter (0.1–100 Hz) and re-referenced to the common average. Two seconds of fade-in and fade-out periods were excluded from the analyses leading to the segmentation of 44s EEG epochs (corresponding to the display of 440 stimuli). The four repetitions were averaged per condition and per participant. A FFT was applied on the signal to extract amplitude spectra for the 64 channels with a frequency resolution (the size of the frequency bins) of 0.023 Hz. We then computed two measures to determine whether and how the brain specifically responded to the deviant frequency in each condition: sum of baseline-corrected amplitudes (SBA), and Z-scores. For the SBA, we computed the baseline-corrected amplitudes by subtracting from the bin of interest (i.e., 1.25 Hz) the mean amplitude of 20 surrounding frequency bins (excluding the one immediately adjacent to the bin of interest), which constitutes the baseline amplitude in microvolts. We calculated similarly the baseline-corrected amplitude of the harmonics of the frequency of interest relative to their neighbors (i.e., up the 7^th^ harmonic of the oddball frequency and up to the 2^nd^ harmonic of the base frequency). Then, we summed the baseline-corrected amplitudes obtained for the frequency of interest and its harmonics and thus obtained a measure that quantifies changes related to the frequencies of interest within the EEG signal. For the z-scores, we cropped the FFT spectra around the frequency of interest (1.25 Hz) and its subsequent harmonics up to the seventh (*i.e.*, 1.25, 2.5, 3.75, 5, 6.25, 7.5, and 8.75 Hz) surrounded by their twenty respective neighboring bins (ten on each side). We summed all cropped spectra and then applied a Z-transformation to the amplitudes. We finally extracted from this Z-transformation the value of the merged critical bin of interest. This value represents the brain response specific to the experimental manipulation at 1.25 Hz (and its harmonics). This score can be interpreted as the strength of the neural detection of the change and assesses the statistical significance of the brain responses to each category change. As a Z-score, a value larger than the value corresponding to the 95th percentile indicates a significant response to the change (*p* < 0.05, one-tailed, testing signal level > noise level). Based on previous results in adults^23^, we regrouped the averaged signal of several electrodes into three “regions of interest” for the analysis: left occipito-parietal (PO7, P7) and right occipito-parietal (PO8, P8) regions of interest; and a bilateral one encompassing the average of both sides to account for lateralization differences between participants.

## Acknowledgment

This research was funded by the European Union’s Horizon 2020 research and innovation program under the Marie Skłodowska-Curie Grant 799171. A.V. is a research associate at the Fonds National de la Recherche Scientifique (FRS-FNRS, Belgium). We thank the master students who recruited the participants and collected the data. APA ethical standards were followed in the conduct of this work.

